# MgrB dependent colistin resistance in *Klebsiella pneumoniae* is associated with an increase in host-to-host transmission

**DOI:** 10.1101/2021.12.01.470879

**Authors:** Andrew S. Bray, Richard D. Smith, Andrew W. Hudson, Giovanna E. Hernandez, Taylor M. Young, Robert K. Ernst, M. Ammar Zafar

## Abstract

Due to its high transmissibility, *Klebsiella pneumoniae* (*Kpn*) is one of the leading causes of nosocomial infections. Here, we studied the biological cost of colistin resistance, an antibiotic of last resort, of this opportunistic pathogen using a murine model of gut colonization and transmission. Colistin resistance in *Kpn* is commonly the result of inactivation of the small regulatory protein MgrB. Without a functional MgrB, the two-component system PhoPQ is constitutively active, leading to increased lipid A modifications and subsequent colistin resistance. Using an engineered MgrB mutant, we observed that MgrB-dependent colistin resistance is not associated with a fitness defect during *in vitro* growth conditions. However, colistin-resistant *Kpn* colonizes the murine gut poorly, which may be due to the decreased production of capsular polysaccharide by the mutant. The colistin-resistant mutant of *Kpn* had increased survival outside the host when compared to the parental colistin-sensitive strain. We attribute this enhanced survivability to dysregulation of the PhoPQ two-component system and accumulation of the master stress regulator RpoS. The enhanced survival of the colistin resistant strain may be a key factor in the observed rapid host-to-host transmission in our model. Together, our data demonstrate that colistin-resistant *Kpn* experiences a biological cost in gastrointestinal colonization. However, this cost is mitigated by enhanced survival outside the host, increasing the risk of transmission. Additionally, it underscores the importance of considering the entire life cycle of a pathogen to truly determine the biological cost associated with antibiotic resistance.

**Importance:** The biological cost associated with colistin resistance in *Klebsiella pneumoniae* (*Kpn*) was examined using a murine model of *Kpn* gut colonization and fecal-oral transmission. A common mutation resulting in colistin resistance in *Kpn* is a loss-of-function mutation of the small regulatory protein MgrB that regulates the two-component system PhoPQ. Even though colistin resistance in *Kpn* comes with a fitness defect in gut colonization, it increases bacterial survival outside the host enabling it to more effectively transmit to a new host. The enhanced survival is dependent upon the accumulation of RpoS and dysregulation of the PhoPQ. Hence, our study expands our understanding of the underlying molecular mechanism contributing to the transmission of colistin-resistant *Kpn*.

## Introduction

Host-to-host transmission of antimicrobial resistant bacteria resulting in nosocomial infections is a significant public health concern and associated with hundreds of billions of dollars in healthcare costs [1, 2]. At the forefront of this threat is the Gram-negative pathogen *Klebsiella pneumoniae* (*Kpn*), which is responsible for the second highest frequency of hospital acquired infections [3]. Carbapenem and extended spectrum β-lactam resistant *Kpn* are associated with poor patient outcomes [4, 5]. The rise in infections due to multi-drug resistant (MDR) *Kpn* has resulted in a correlative increase in the use of last-resort antibiotics, such as polymyxins [6, 7]. Polymyxins are cationic anti-microbial peptides (cAMPs) that disrupt the outer membranes of Gram-negative bacteria by interacting with the negatively charged lipopolysaccharides (LPS) on the bacterial membrane surface [8]. Unfortunately, increasing use of polymyxins has fueled the generation of polymyxin resistant *Kpn* strains [9]. While initially reported outbreaks of MDR and polymyxin resistant *Kpn* were largely in South-East Asia, these strains have now spread around the globe [10-13].

Clinically there are two commonly used polymyxins, colistin (polymyxin E) and polymyxin B, which share a similar structure of a cationic peptide ring with a linear chain of three amino acids that connect to a hydrophobic acyl tail [6, 8]. Treatment with colistin is considered a salvage therapy for patients infected with MDR Gram-negative bacteria [6, 7, 14]. Colistin resistance is well characterized in *Kpn* and involves modifications to the negatively charged lipid A moiety of the lipopolysaccharide (LPS) [15] that occur through mutations resulting in dysregulation of the two-component systems (TCS) PmrAB, PhoPQ, or CrrAB [16-18]. PmrAB normally responds to ferric iron and mild acidic conditions which results in PmrB phosphorylating the response regulator PmrA [19, 20]. Phosphorylated PmrA in turn activates expression of the *arnBCADTEF* operon, *ugd*, and *pmrC* [21, 22]. Both the *arn* operon and *ugd* mediate the addition of 4-aminoarabinose to the lipid A moiety whereas PmrC mediates the addition of a phosphoethanolamine [21, 22]. The CrrAB system is poorly characterized. However, phosphorylated CrrA positively regulates the expression of the *arn* operon and *crrC*, which interacts with PmrA resulting in the further addition of phosphoethanolamine [23, 24]. Under cAMP stress or low magnesium environments, PhoQ phosphorylates PhoP, driving PhoP interaction with PmrA as well as directly regulating the *arn* operon [17, 20], thus ensuring the additions of 4-aminoarabinose and phosphoethanolamine to LPS [25, 26]. Additionally, PhoP directly activates the expression of *pagP* and *lpxO*, which are involved in palmitoylation and hydroxylation of the lipid A moiety, respectively [27, 28]. More recently, colistin resistant strains have been identified carrying *mcr-1* on a plasmid, the product of this gene is a transferase that directly adds phosphoethanolamine to the lipid A moiety [29].

Despite the contribution of the multiple TCSs in lipid A modification, colistin resistance in the majority of *Kpn* strains is caused by mutations that abrogate the function of the small regulatory protein MgrB [30, 31]. MgrB, localizes to the inner membrane and negatively regulates PhoPQ activity. Consequently, PhoPQ is over-activated when MgrB is inactivated, resulting in an increase in the lipid A modifications that confer colistin resistance [17, 32]. Inactivation or gain of function mutations in PhoP, PhoQ, PmrA, PmrB, or CrrB are also known to result in colistin resistance, but MgrB inactivation remains the most prevalent cause of resistance in *Kpn* [16-18, 23, 24, 30].

Anti-microbial resistance targets specific biological functions of the cell and consequently it is generally associated with a biological cost. The biological cost of antibiotic resistance can manifest as altered bacterial growth, virulence, or transmissibility [33]. The cost of colistin resistance has been studied in a variety of *Enterobacteriaceae*, including *Kpn*, with conflicting results [34-37]. The inconsistent data on biological cost can be attributed to the type of mutation, the strain tested, and on the type of assays (*in vivo* and *in vitro)* [34-37]. However, studies investigating the biological cost of colistin resistance in *Kpn* have neglected the potential impact on the common initial colonization step-the gastrointestinal tract (GI) -as well as host-to-host transmission [36, 37]. Epidemiological data suggest the GI tract to be the initial site of colonization for *Kpn*, from where it can cause disease in the same individual or transmit to another host [38]. Our recent murine model of *Kpn* GI colonization with an intact microbiome experimentally confirmed that *Kpn* colonizes the GI tract, and transmits to another host through the fecal-oral route [39]. Furthermore, in our murine model we observed that the *Kpn* MDR isolate (KPNIH1) poorly colonized the murine GI tract compared to non-MDR isolates, suggesting a possible biological cost associated with antibiotic-resistance, though the KPNIH1 strain is genetically distinct in multiple ways from the KPPR1S strain used in this publication [39]. The importance of understanding the initial GI colonization by *Kpn* is two-fold: first, *Kpn* gut colonization is associated with poor patient outcome [38, 40], and second, *Kpn* is generally considered a silent gut colonizer and thus transmits asymptomatically, posing a major threat in hospital settings where multiple potential reservoirs could be silently transmitting and making it difficult to identify and control the spread of infection [39, 41].

Given the increased frequency of resistance to colistin, an antibiotic of last resort, we assessed the potential biological cost of MgrB dependent colistin resistance by using our murine model that allows us to follow the entire lifecycle of *Kpn*. Ultimately this may aid in devising better strategies to combat the spread of pathogenic microbes in a hospital setting.

## Results

### MgrB dependent colistin resistance of *Kpn* does not affect *in vitro* growth but provides enhanced survival against cAMP

We first examined the biological cost of MgrB dependent colistin resistance through growth assays using nutrient rich (Lysogeny Broth [LB]) media and minimal media (M63). We compared KPPR1S (a derivative of ATCC43816; WT; AZ55, **Table S1**) to its isogenic mutant *ΔmgrB* (MgrB^-^; AZ132, **Table S1**), and the chromosomally complemented (reconstituted at the native site) strain (MgrB^+^; AZ141, **Table S1**). All three strains grew equally well with no differences in generation time in both media conditions, suggesting the MgrB^-^ strain has no fitness defect under *in vitro* growth conditions (**Fig S1*A-B***).

Abrogated function of MgrB results in modifications of the lipid A domain of LPS, allowing for protection against the cation mediated targeting and disruption of the bacterial cell membrane by colistin [30, 31]. In order to determine the potential impact of MgrB-dependent colistin resistance on survivability against other cAMPs we tested the survival of the WT, MgrB^-^, and MgrB^+^ strains against lysozyme, as it is one of the most abundant enzymes in the mucosal surface and has cAMP activity [42], and polymyxin B, a cAMP similar to colistin. MgrB^-^ had higher survival rates, as compared to the parental WT strain when treated with increasing concentrations of either lysozyme or polymyxin B (**Fig. 1*A-B***). The MgrB^+^ strain behaved similarly to WT, suggesting that the enhanced survival of the MgrB^-^ strain is specific to inactivation of MgrB. Taken together, our data show that MgrB dependent cAMP resistance is not associated with reduced growth *in vitro*.

**Fig 1.**
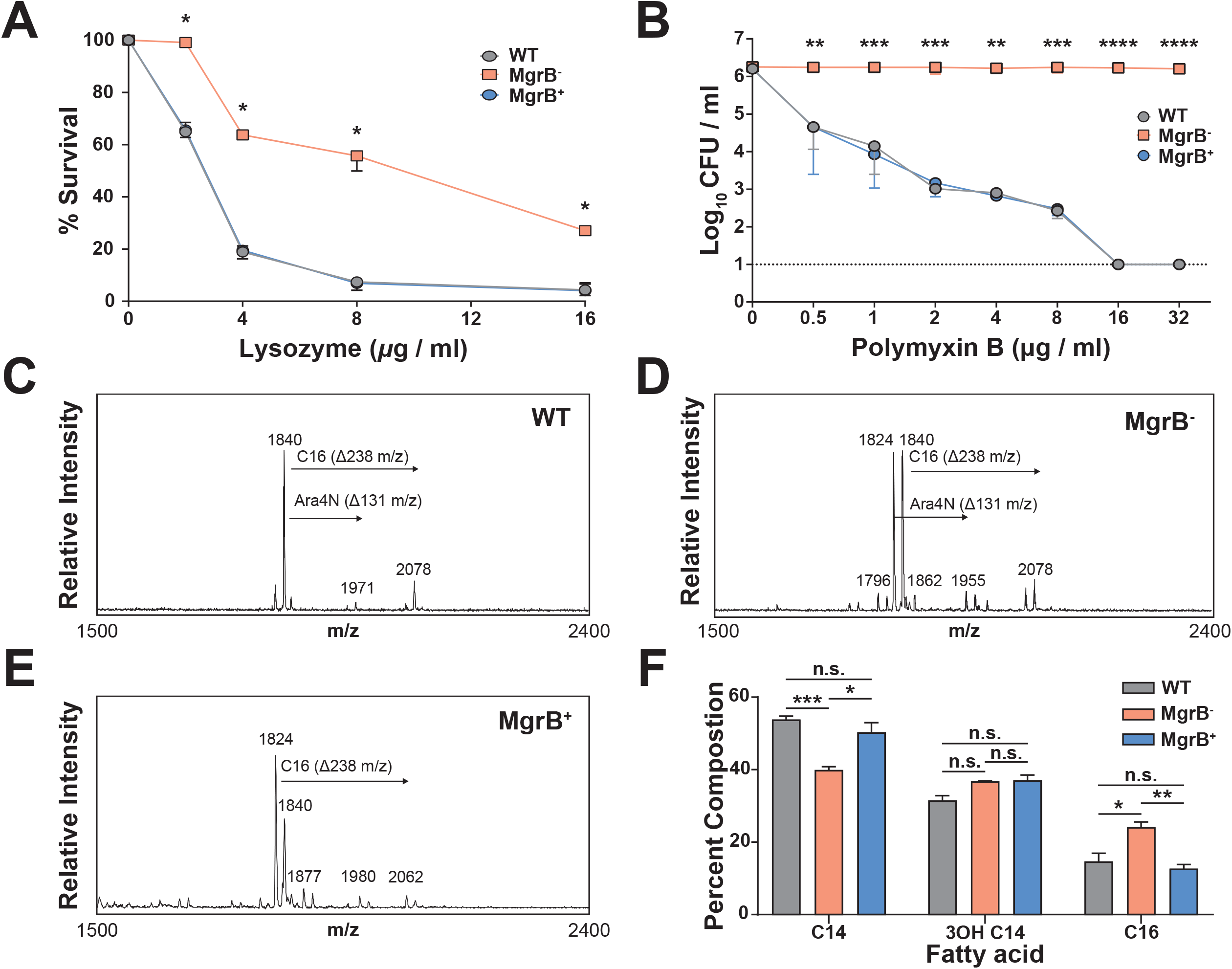
MgrB dependent colistin resistance provides enhanced survival against cAMP. *Kpn* survival against (***A***) lysozyme and (***B***) polymyxin B. (***A***) Mid-log phase (OD_600_) cultures of WT, MgrB^+^, and MgrB^-^ were re-suspended in PBS and incubated with the appropriate concentrations of lysozyme for 1 hour at 37°C. (***B***) Mid-log phase (OD_600_) cultures of each strain were diluted to ∼10^5^ CFU / ml in LB before being incubated with increasing concentrations of polymyxin B for 30 minutes at 37°C. Fatty acid composition was conducted using GC/MS via the fatty acid methyl ester (FAME) derivatization from bacterial culture pellet. Strains were analyzed with 3 biological replicates for ***(C)*** WT, ***(D)*** MgrB^+^, and ***(E)*** MgrB^-^. (***F***) Fatty acid composition of lipid A of each *Kpn* strain. Statistical differences were calculated using Kruskal-Wallis tests with Dunn’s test of multiple comparisons within each concentration group (***A-B***) or fatty acid group (***F***). There was no significant difference between the WT and MgrB^+^ strains. * *P* < 0.05, ** *P* < 0.005, *** *P* < 0.0005, **** *P* < 0.00005

The fatty acid composition of the lipid A for WT, MgrB^-^, and MgrB^+^ were evaluated using gas chromatography (GC) (**Fig. 1*C-E***). As shown in **Fig. 1*F***, the lipid A for all three strains was composed of C14, 3OH-C14, and C16 acyl groups. WT and MgrB^+^ lipid A had similar fatty acid composition, whereas MgrB^-^ had significantly lower proportion of C14 and higher proportion of C16. The difference in C14 and C16 concentrations may be explained by potential increase in PagP activity, which adds the C16 palmitoyl modification to lipid A, which is upregulated by PhoP [27]. Additionally, it has been shown that PhoPQ has a down-regulatory effect on the acyltransferases LpxL1 and LpxL2 which mediate C12 and C14 additions to lipid A, respectively [43]. Furthermore, the increased relative abundance of C16 is one potential mechanism for conference of colistin resistance as it alters the rigidity of the outer membrane [44].

### MgrB^-^ *Kpn* has a fitness defect in gut colonization that is rescued by antibiotic treatment

As antibiotic resistance can be associated with biological costs at various stages of a pathogens life cycle (colonization, infection, transmission) we decided to use our published animal model to test the ability of the MgrB^-^ strain to colonize adult, immunocompetent, non-antibiotic treated mice [39]. We inoculated mice with either WT, MgrB^-^, or MgrB^+^ strains of *Kpn* and enumerated bacterial shedding from their feces. Over the course of 15 days of infection, the MgrB^-^ strain shed poorly in comparison to the parental WT strain (**Fig. 2*A***), indicating a potential biological cost of MgrB-dependent colistin resistance in *Kpn* may manifest as a defect in gut colonization. The MgrB^+^ strain colonized as well as WT, suggesting that the colonization defect is specific to inactivation of MgrB. To ensure that the observed colonization defect of MgrB-is not strain specific, we infected mice with a clinical fecal isolate and its isogenic *mgrB* mutant (ST1322 and ST1322 (MgrB^-^), AZ99 and AZ179, **Table S1**). We had previously shown that ST1322 colonizes the murine GI tract, albeit at a lower density [39]. Compared to ST1322 the ST1322 (MgrB-) colonizes poorly, and is cleared by the host 3 days post-infection, suggesting that the poor colonizing ability of the MgrB-is not strain specific (**Fig. S2**).

**Fig 2.**
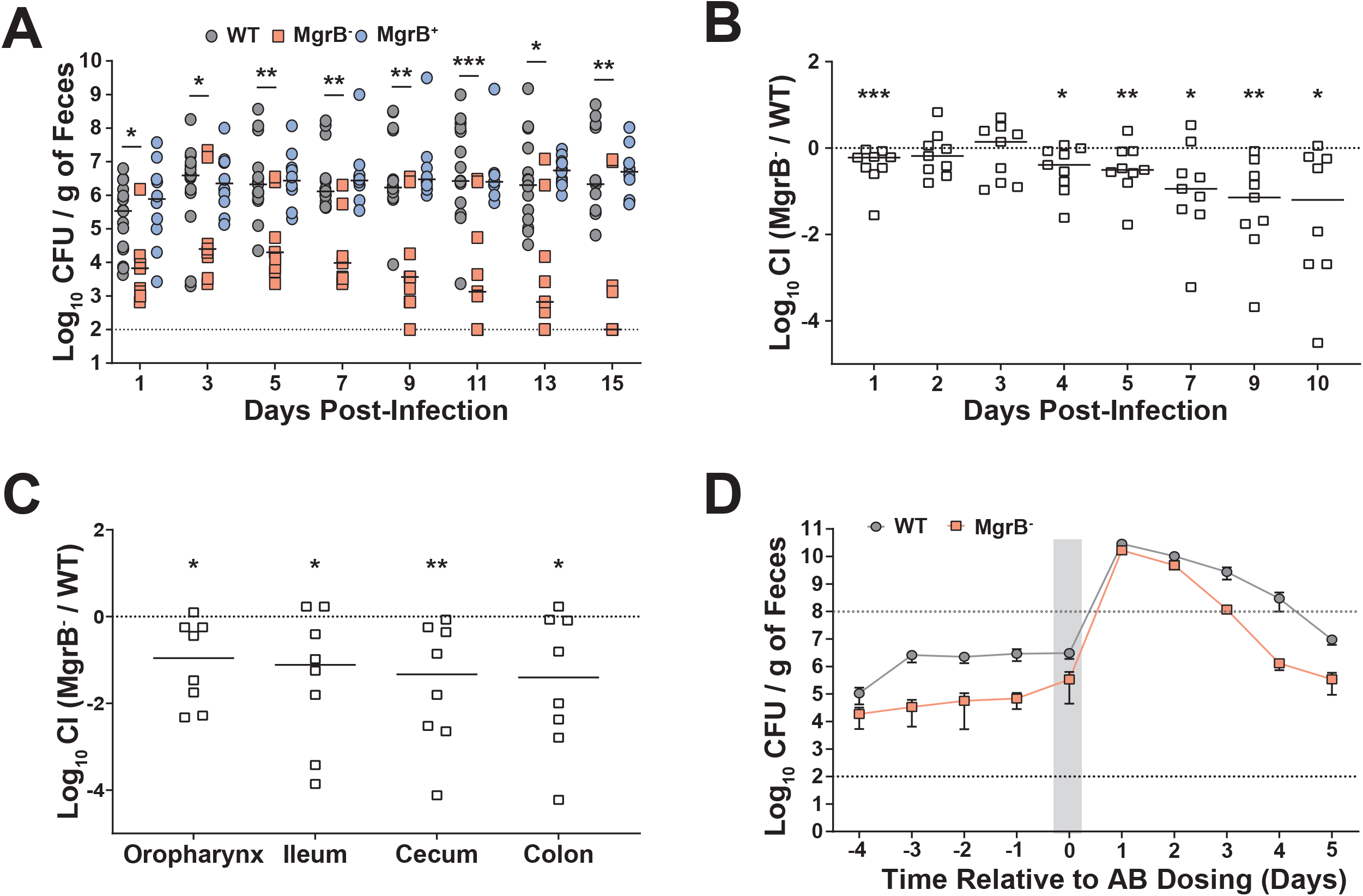
MgrB^-^ has a fitness defect in gut colonization that is rescued by antibiotic treatment. (***A***) Fecal shedding of infected mice. Mice were orally infected with either WT, MgrB^-^, or MgrB^+^ *Kpn*, and feces was collected on the days indicated (n ≥ 10 for each group). Each point indicates a single mouse on a given day, the bars indicate the median shedding, and the dotted line indicates the limit of detection, significance shown is between WT and MgrB^-^. Kruskal-Wallis followed by Dunn’s test of multiple comparisons was performed at each time point for analysis, WT and MgrB^+^ were not significantly different. (***B***) Competitive index (CI) of infected mice. Mice were orally infected with a 1:1 mixture of the WT and MgrB^-^ mutant, with feces collected on the days indicated (n ≥ 10). The CI was determined as described in Materials and Methods. Each symbol represents the log_10_ CI value from an individual mouse on a given day, with a bar indicating the median value. The dashed line indicated a competitive index of 1 or a 1:1 ratio of WT to mutant. (***C***) Colonization density of the colon, cecum, ileum, and oropharynx represented as log_10_ CI value from an individual mouse at the end of the study. (***D****) Kpn* Fecal shedding from mice treated with antibiotic (AB). Mice were orally infected with either WT or the MgrB^-^, and 5-days post-infection were given a dose of 5 mg of streptomycin via oral gavage. The shaded gray area indicates the duration of antibiotic treatment, shown are the means and standard error of the means for both WT (*n* = 6) and MgrB^-^ (*n* = 6) infected mice, the black dotted line indicates the limit of detection, the gray dotted line indicates the super shedder threshold. For CI, statistical differences were determined by Wilcoxon signed-rank test. * *P* < 0.05, ** *P* < 0.005, *** *P* < 0.0005

We next assessed whether WT *Kpn* could compensate *in trans* for the observed defect in GI colonization by MgrB^-^ during co-infection. Co-infected mice still displayed poor shedding and colonization of MgrB^-^ *Kpn* compared to WT (**Fig. 2B-C**). These observations suggest that the parental WT strain is unable to rescue the defect in gut colonization observed from the MgrB^-^ strain. A single dose of antibiotics causes the development of a super-shedder state (>10^8^ CFU/g of feces), where the amount of *Kpn* shed from antibiotic treated mice is ∼100 fold higher when compared to mock treated mice [39]. We hypothesized that reducing the gut flora through antibiotic treatment would allow MgrB^-^ *Kpn* to colonize better than in the presence of normal gut flora. Mice infected with either the WT or MgrB^-^ were given a single dose of streptomycin 5 days post-infection. We found that antibiotic treatment triggered a temporary super-shedder phenotype in both the WT and the MgrB^-^ strains, with both reaching the same high density in the feces (**Fig. 2*D***). However, post-treatment the WT strain continued to shed at a higher level longer when compared to the MgrB^-^ strain, suggesting that continuous treatment of antibiotic is required to bypass the intrinsic colonization defect of the MgrB^-^ strain. Together these results show that there is a biological cost associated with inactivation of MgrB in *Kpn* in the context of GI colonization, and that antibiotic treatment alleviates this fitness defect.

### Loss of functional MgrB impacts *Kpn* capsular polysaccharide production and its interaction with mucus

*Choi et al*. observed variability in production of biofilm, capsular polysaccharide (CPS), and hypermucoviscosity (HMV) between spontaneously generated colistin resistant mutants of *Kpn* and their respective parental strains [37]. Therefore, we assessed whether MgrB inactivation would have pleiotropic effects impacting these three *Kpn* virulence phenotypes. No differences were observed in biofilm formation or the HMV phenotype between the WT and MgrB^-^ strains (**Fig. S3*A-B***). To assess whether the WT and mutant differed in level of CPS, we quantified the amount of uronic acid, the major component of *Kpn* CPS produced by these strains. Compared to the WT strain, MgrB^-^ produced less CPS (**Fig. 3*A***). We have shown previously that CPS is required for robust *Kpn* GI colonization [39]. Additionally, CPS prevents clearance by allowing bacteria to escape the mucus present on the mucosal epithelial layer [45]. Thus, we hypothesized that reduced CPS in the MgrB^-^ strain leads to entrapment in the mucus layer, eventual clearance, and thus reduced GI colonization. To determine whether reduced capsule amount leads to increased clearance, we carried out an *in vitro* solid-phase mucin binding assay with semi-purified mucin, the principal component of mucus. The un-encapsulated mutant (CPS^-,^ *ΔwcaJ*, AZ124, **Table S1**) showed significantly higher binding to bovine submaxillary mucin compared to the WT strain (**Fig. 3*B***). Moreover, the MgrB^-^ strain with reduced CPS had enhanced binding to mucin compared to the WT and MgrB^+^ strains (**Fig. 3*B***). Taken together, our results suggest a correlation between a reduced ability to bind mucin through the production of CPS and robust GI colonization. This implies that the biological cost of MgrB-dependent colistin resistance is not necessarily associated with a growth defect, but dysregulation of CPS production leading to enhanced clearance.

**Fig 3.**
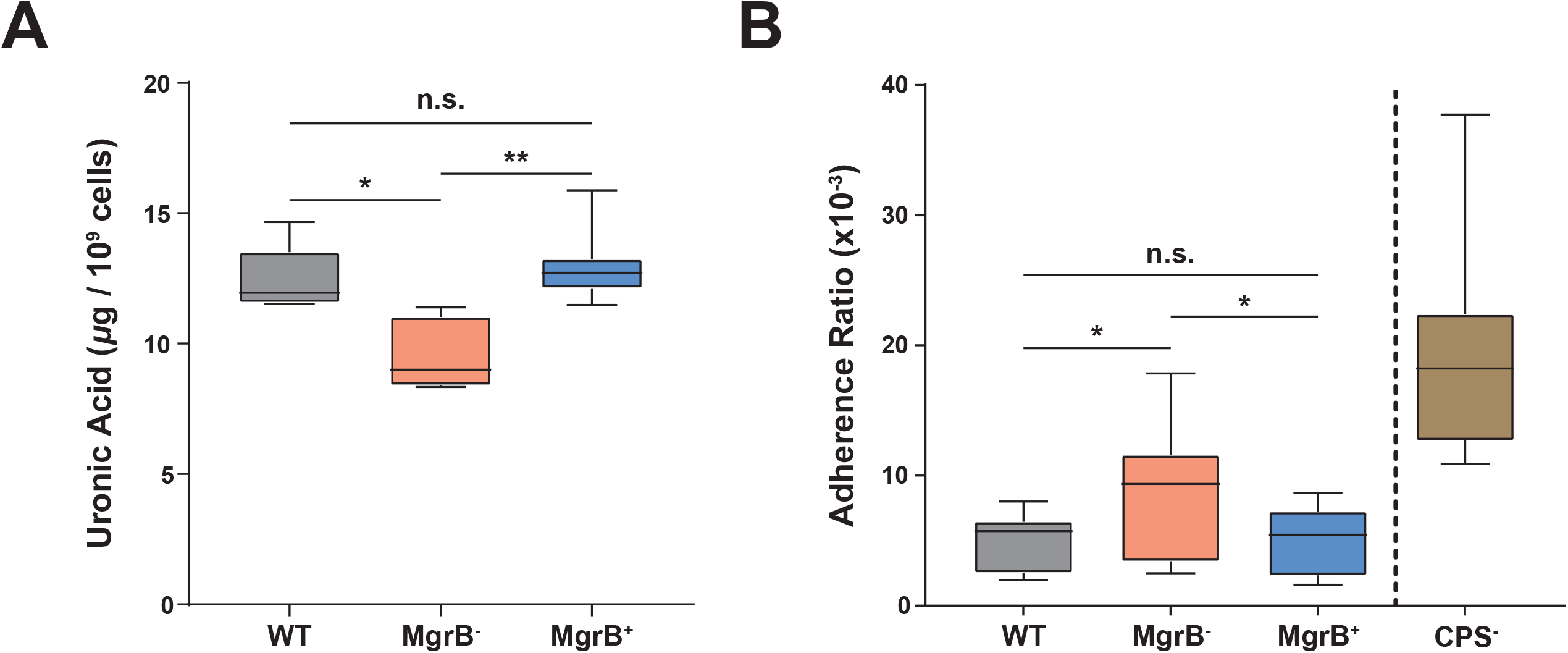
MgrB inactivation reduces capsule amount and increases mucus association. (***A***) Comparison of capsule production between WT, MgrB^-^, and MgrB^+^ using a uronic acid assay from stationary phase culture samples. (***B***) Comparison of the association of WT, MgrB^-^, MgrB^+^, and CPS^-^ (*Δwcaj*) to immobilized semi-purified bovine submaxillary gland mucin. Shown is the adherence ratio 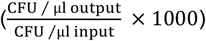 of each strain. Box and whiskers indicate the means and minimum to maximum values, three independent assays were carried out for both capsule amount (in duplicate) and mucin binding (in sextuplicate). Statistical differences were calculated using Kruskal-Wallis tests with Dunn’s test of multiple comparisons across strains. * *P* < 0.05, ** *P* < 0.005, n.s, not significant.

### MgrB inactivation increases *Kpn* survival outside the host

In addition to protecting bacteria from host mediated clearance mechanisms, capsule contributes towards environmental survival [46]. Fomites, environmental reservoirs of infections such as catheters, ventilators, or bed hand-rails, are considered a significant source of *Kpn* transmission in a hospital setting [47], and coupled with our finding that MgrB^-^ has reduced capsule compared to the WT strain, (**Fig. 3*B***) we decided to test the environmental survivability of our strains. To determine survivability, we conducted solid surface starvation survival experiments by drying WT, MgrB^-^, and MgrB^+^ strains onto nitrocellulose membranes and placing them on agarose pads. *Kpn* is unable to metabolize purified agarose and nitrocellulose as energy sources, and the agarose pads allowed us to provide stable temperature and humidity. Bacteria suspended in PBS were spotted on to the agarose pads, with viable count determined over a period of 12 days. CFU’s for the WT and MgrB^+^ declined rapidly over the first few time-points (**Fig. 4*A***). However, MgrB^-^ did not have a sharp decline in viable count and showed a significantly higher survival compared to the WT and the complemented strain (**Fig. 4*A***). Furthermore, we tested the survivability of ST1322 and its isogenic MgrB-mutant. Remarkably the phenotype was conserved, with ST1322 (MgrB^-^) surviving at a significantly higher rate than its parental strain, ST1322 (**Fig. 4*B***).

**Fig 4.**
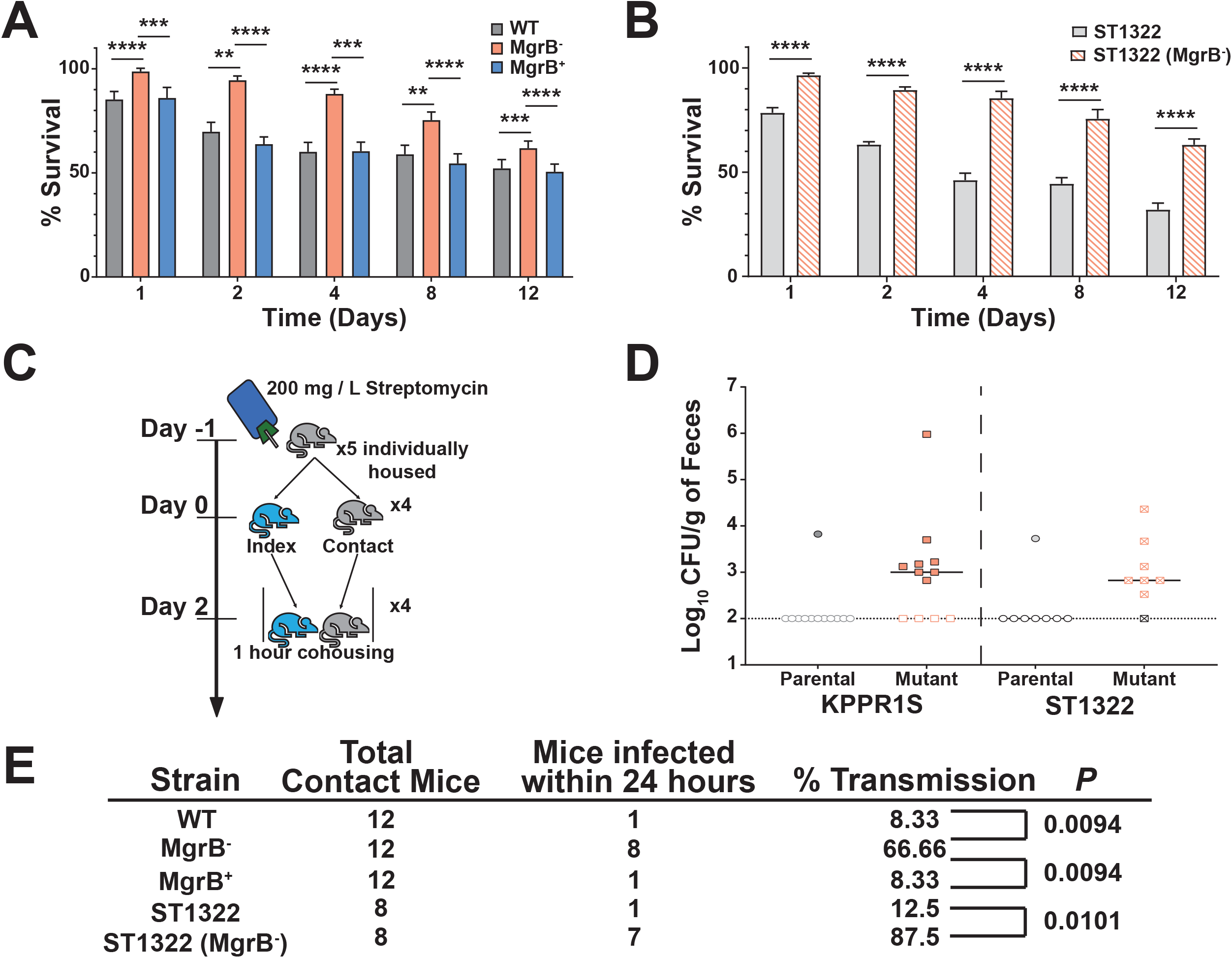
MgrB inactivation affects survival outside the host and enhances host-to-host transmission. (***A-B****) Ex vivo* solid surface survival of *Kpn* isolate KPPR1S and ST1322, their *ΔmgrB* isogenic mutants, and the chromosomal complement *mgrB*^*+*^. Bacterial strains were grown overnight, re-suspended in PBS, OD_600_ adjusted to 4, and spotted on to nitrocellulose discs on 1% agarose pads in 6-well polystyrene plates. Discs were removed at appropriate time interval, and bacteria re-suspended in PBS and viable bacterial counts determined. Shown is the mean and SEM. Statistical analysis was carried out using Kruskal-Wallis tests with Dunn’s test of multiple comparisons at each time point (***A***) or using Mann-Whitney U tests at each time point (***B***). * *P* < 0.05, ** *P* < 0.005, *** *P* < 0.0005, **** *P* < 0.00005, n.s, not significant. (***C***) Schematic representation of the protocol for the transmission studies. Mice were placed on antibiotic water 24 hours before separating and infecting the index mouse. Once the index mouse was robustly colonized, the individually housed contact mice were exposed to the index mouse for 1 hour. Fecal shedding was collected the following day to determine if transmission had occurred. (***D***) Fecal shedding of *Kpn* parental isolates (KPPR1S and ST1322) and their isogenic mutant (*ΔmgrB*) from contact mice after a single exposure, the bar indicates the median shedding, dotted line indicates limit of detection. (***E***) Summary of transmission data for the indicated strains after a single 1-hour exposure. The *p* value was calculated using the Fisher’s exact test.

As fomites are a significant source of transmission in a hospital setting [47], we hypothesized that augmented environmental survival of MgrB^-^ could manifest as enhanced host-to-host transmission. Therefore, we assessed the transmission efficiency of the MgrB^-^ and ST1322 (MgrB^-^) strains compared to the WT and ST1322 strains, respectively. As MgrB^-^ colonizes the GI tract poorly, but reaches similar levels of colonization to WT under antibiotic pressure (**Fig. 2D**), mice were individually housed with water containing streptomycin. One mouse (Index) was infected with WT, MgrB^-^, ST1322, or the ST1322 (MgrB^-^) strain and the uninfected mice (contact) were subsequently exposed to the index mouse in the infected mouse’s cage for 1 hour each day for up to 5 days (**Fig. 4*C***). Transmission was assessed by determining if contact mice fecal pellets were positive for *Kpn* (shedding). All strains colonized index mice at high density (**Fig. S4*A***), with the MgrB^-^ index mice colonized at a slightly reduced level. After a single one-hour exposure more than 60% of the MgrB^-^ contact mice were colonized with MgrB^-^ *Kpn*, whereas less than 10% of the WT contact mice were colonized (**Fig. 4*D-E***). Similarly, we observed higher transmission post one-hour exposure with the ST1322 (MgrB^-^) when compared to its parental strain (80% vs 12.5%) (**Fig. 4*D-E***). Thus, increased environmental survival correlates with higher transmission efficiency, which is not strain specific. Furthermore, once colonized the contact mice remained colonized for the duration of the study (**Fig. S4*B***).

### Mechanism for increased environmental survival

MgrB is known to have regulatory function in two key molecular pathways, repression of PhoPQ and sequestration of the small protein IraM. IraM is an anti-adaptor of SprE/RssB, the protein that chaperones the stress response master regulator, RpoS, to the ClpXP protease for degradation [48]. In *mgrB* mutants of *Escherichia coli* there is stabilization and subsequent accumulation of RpoS due to the negative regulation of SprE/RssB by IraM [48]. We observed an accumulation of RpoS in the MgrB^-^ strain (**Fig. S5**), and hypothesized that an increase in RpoS, would be the key factor increasing MgrB^-^ *Kpn* survival under the stress induced conditions of solid surface starvation. Thus, a *rpoS* deletion mutant (RpoS^-^, AZ139, **Table S1**) survived poorly when compared to the WT strain (**Fig. 5*A***). However, a *mgrB, rpoS* double deletion mutant strain (MgrB^-^/RpoS^-^, AZ138, **Table S1**) partially restores survivability when compared to the RpoS^-^ strain, suggesting that although RpoS promotes environmental survival there is an additional mechanism(s) that impacts *Kpn* environmental survival (**Fig. 5*A***). Next, we determined the contribution of PhoPQ to environmental survival of *Kpn*. A *phoQ* deletion mutant (PhoQ^-^, AZ107, **Table S1**) had lower CFU’s compared to the parental WT strain, especially at Day 1 (**Fig. 5*B***). Additionally, a double mutant that inactivated *mgrB* and *phoQ* (MgrB^-^/PhoQ^-^, AZ150, **Table S1**) had similar survival to the WT isolate, indicating that PhoQ over-activation in the absence of MgrB is a significant contributor to solid surface starvation survival of *Kpn*.

**Fig 5.**
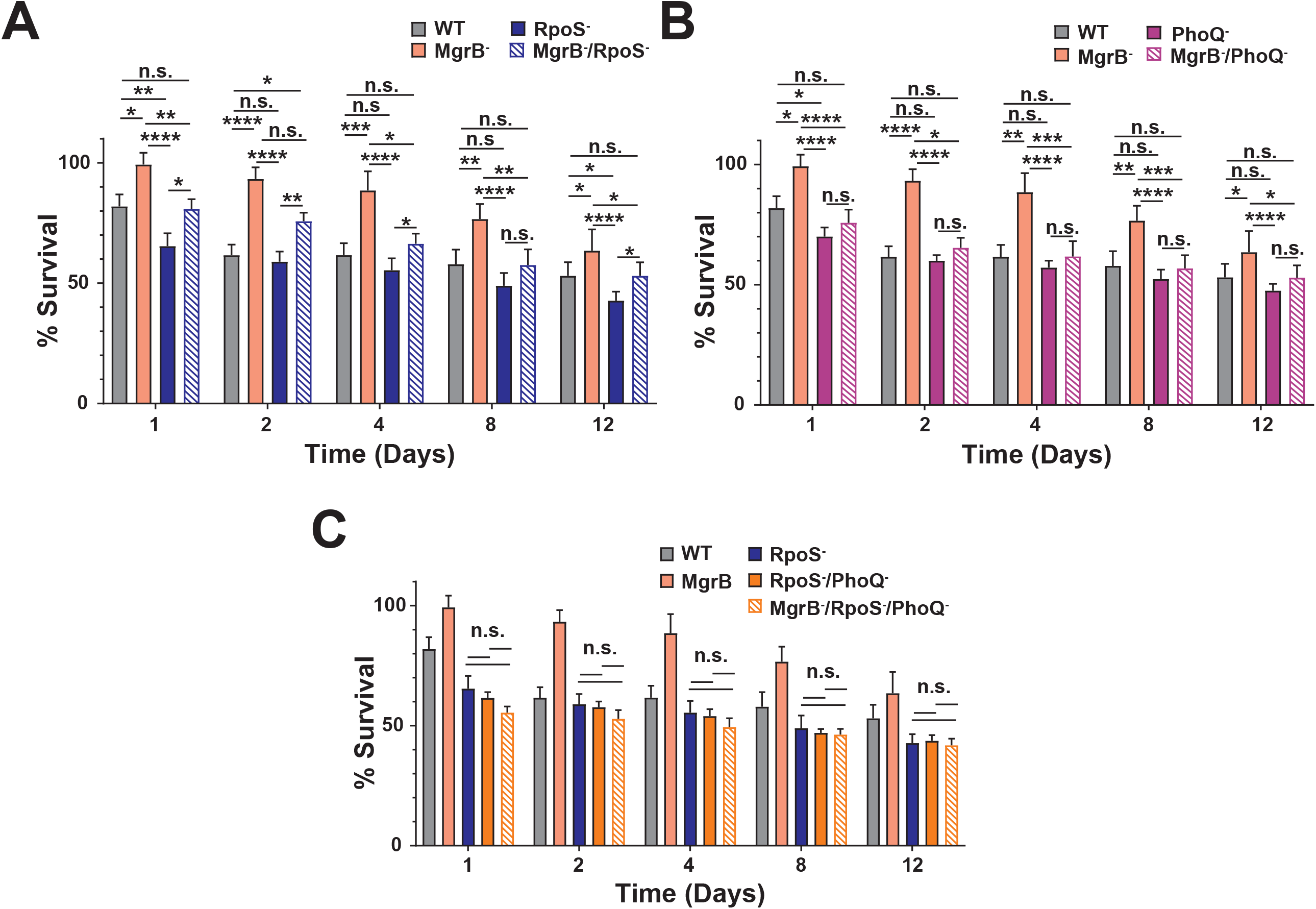
Contribution of TCS PhoPQ and the master stress regulator RpoS to the environmental survival of *Kpn. Ex vivo* solid surface survival of KPPR1S and its isogenic mutants. (***A***) The impact of loss of function mutations in MgrB-, RpoS- and the double mutant MgrB-, RpoS-on *Kpn* survival. (***B***) The contribution of the sensor kinase PhoQ in the environmental survival of *Kpn*. (***C***) MgrB-dependent enhanced survival is dependent upon both RpoS and PhoQ. The WT and MgrB^-^ data is repeated in each panel. Statistical differences were calculated using Kruskal-Wallis tests with Dunn’s test of multiple comparisons across strains at each time point. * *P* < 0.05, ** *P* < 0.005, n.s, not significant.

Furthermore, we also determined whether a *rpoS* and *phoQ* double deletion mutant (RpoS^-^ /PhoQ^-^, AZ151, **Table S1**) would have an additive effect on reduced survivability, with and without *mgrB* inactivation (MgrB^-^/RpoS^-^/PhoQ^-^, AZ155, **Table S1**). The double mutant (RpoS^-^ /PhoQ^-^) does have reduced survivability compared to the WT and the individual mutants (**Fig. 5*C***). Moreover, the triple mutant (MgrB^-^/RpoS^-^/PhoQ^-^) behaves similarly to the double mutant (RpoS^-^/PhoQ^-^), with the MgrB inactivation unable to rescue the defect, suggesting that the increase in environmental survival through MgrB inactivation is dependent on its dysregulation of PhoPQ and RpoS.

## Discussion

This study provides a comprehensive understanding of MgrB dependent colistin resistance as it pertains to the ability of *Kpn* to colonize, survive, and transmit from host to host. We show that the loss of functional MgrB plays a role throughout the pathogenic life-cycle of *Kpn*. The dysregulation caused by inactive MgrB attenuates *Kpn* gut colonization, but increases survival outside the host which results in more rapid transmission. This new understanding of the effects of Colistin resistance on infection and transmission of *Kpn* may guide future investigations into how other antibiotic resistances impact bacteria, as well as re-examine the quarantine and containment practices of patients infected with such pathogens

While animal models have allowed for the study of *Kpn* lung colonization [49, 50], the initial colonization of the gut by *Kpn* is poorly understood. Historically this is due to ineffective *in vivo* models that utilized antibiotic treatment and infection via oral gavage to establish colonization [51, 52]. Using our recently established murine model [39] we assessed the ability of colistin resistant *Kpn* to establish itself in healthy, immunocompetent mice with intact microbiota. Our data show that colistin resistant *Kpn* (MgrB^-^) has a fitness defect in GI colonization (**Fig. 2*A***). While this result is promising for the implications of reduced asymptomatic carriage and shedding, we also observed an increase in survival against the host defense mechanism of lysozyme (**Fig. 1*B***) and recovery of shedding and colonization to WT levels under antibiotic treatment (**Fig. 2*D***). Moreover, we examined several known virulence phenotypes of *Kpn* to identify the underlying mechanism for the observed fitness defect in gut colonization by the MgrB^-^ strain. We focused on biofilm formation, hypermucoviscosity, and capsule production, as they are energy dependent processes and previously implicated with an antibiotic resistant dependent biological cost [37]. We observed reduced capsule production by the MgrB^-^ strain that correlates with increased mucin binding (**Fig. 3**), indicating that the observed fitness defect in gut colonization could be due to an increase in mucosal clearance of MgrB^-^ *Kpn*.

Our model also allowed us to study another poorly understood facet of *Kpn* infection, host-to-host transmission [39]. *Kpn* readily transmits through the fecal-oral route and in a hospital setting has been associated with several super-spreading events [40] primarily through fomites [47] making it difficult to track outbreaks. Moreover, *Klebsiella* species can survive on various surfaces for extensive periods of time, which allows them ample opportunity to be acquired by another susceptible individual [47, 53]. Conversely, environmental survivability of specific *Kpn* mutants is greatly under-studied. Here we show that the MgrB^-^ strain surprisingly had enhanced survivability compared to the WT (**Fig. 4*A***), and that this phenotype was not strain specific (**Fig. 4*B***). With the majority of colistin-resistant strains of *Kpn* linked to inactivation of the small regulatory protein MgrB [30, 31], it seems that this particular genetic variation would not contribute significantly to environmental survival. However, in addition to regulating PhoPQ, which has a myriad of effects on LPS and the outer membrane [17, 20, 25-28], MgrB also sequesters IraM, which functions as an anti-adaptor to the chaperone protein SprE that binds and delivers the stress response master regulator RpoS for proteolytic degradation [17, 32, 48]. Additionally, phosphorylated PhoP upregulates the transcription of IraP, another anti-adaptor of SprE [54]. Therefore, the consequences of inactivation of MgrB reach beyond colistin resistance and can result in outer-membrane alteration and a constitutive stress response from the bacteria. Both responses potentially contribute to the environmental survival and subsequent transmission of *Kpn*.

We initially hypothesized that the observed increase in survival would be due to a priming effect as a result of accumulating RpoS. Surprisingly, even though RpoS levels were higher in our MgrB^-^ *Kpn* strain, this increase only led to a minor contribution in environmental survival (**Fig. 4*C***), instead the effect is mainly attributable to the PhoPQ two-component system (**Fig. 4*B***). It is likely that the lipid A modifications observed in MgrB-inactivated *Kpn* linked to colistin resistance are also responsible for the increased environmental survival, as lipid A modifications have been previously linked to membrane stabilization and increased survival [55, 56]. While MDR is associated with a biological cost that can manifest itself in colonization and affect transmissibility in a hospital setting (fitness cost) [33], studies ascertaining a biological cost associated with antibiotic resistance generally do not focus on host-to-host transmission.

Herein, we showed that colistin resistant *Kpn* under oral antibiotic pressure colonized as well as the parental strain (**Fig. 2*D***), but transmitted at a higher frequency (**Fig. 4*D***). These results suggest a gain of function. Furthermore, a recent molecular epidemiological study by *Lapp et al* suggests that certain colistin-resistant *Kpn* isolates of the ST258 lineage have enhanced transmissibility attributed to the disruption of the putative O-antigen glycosyltransferase *kfoC* [57]. While *Kpn* is highly genetically heterogeneous, our transmission data with KPPR1S and ST1322 and their respective *ΔmgrB* isogenic mutants are in line with their data [57]. The observed increased transmissibility in our isolates is likely due to the enhanced survivability of the MgrB mutant strain in the environment compared to their parental strains (**Fig. 4A-B**).

Taken together, our study experimentally advances our understanding of the cost associated with MgrB dependent colistin resistance, and underscores the importance of considering the full life cycle of the pathogen when characterizing the biological cost associated with antibiotic resistance. While MgrB^-^ *Kpn* may colonize healthy individuals poorly, those on antibiotic treatment can become robustly colonized and transmit rapidly. Even in non-antibiotic treated individuals who likely experience poor colonization, there is the concerning potential for increased horizontal gene transfer of antibiotic resistances, a problem that has already been linked to *Kpn* and could grow with increased environmental survival and transmission [58, 59]. Colistin resistant isolates can spread even with appropriate infection control practices. Thus, our study underscores the need for an improved antimicrobial stewardship as well as preventative sterilization methods to curb the potential increase in transmissibility of colistin resistant *Kpn*.

## Acknowledgements

We are grateful for the members of Zafar (Wake Forest School of Medicine), Ernst (UMB) and Miller (UNC Chapel Hill) laboratories for fruitful discussions and comments on the manuscript. This project was supported by startup funds provided by Wake Forest Baptist Medical Center to M.A.Z. A.S.B was supported by the NIAID training program in immunology and pathogenesis (5T32AI007401-28). RKE and RDS were supported by the NIH (R01AI104895).

## Methods

### Ethics Statement

This study was conducted according to the guidelines outlined by National Science Foundation animal welfare requirements and the *Public Health Service Policy on Humane Care and Use of Laboratory Animals* [60]. All animal work was done according to the guidelines provided by the American Association for Laboratory Animal Science (AALAS) [61] and with the approval of the Wake Forest Baptist Medical Center Institutional Animal Care and Use Committee (IACUC).

The approved protocol number for this project is A20-084.

### Strain construction

Strains, plasmids and primers used used in this study are listed in **Tables S1, S2** and **S3** respectively. The deletion mutant (*ΔmgrB*, AZ132) was constructed as described [62]. Briefly, a kanamycin cassette bookended with FRT sites from plasmid pKD4 with 60-bp homology to the upstream and downstream region of *mgrB* was PCR amplified using high fidelity Q5 polymerase (New England BioLabs [NEB]). The purified PCR product was then electroporated into the target strain (AZ63) containing the temperature sensitive plasmid pKD46 (**Table S2**), containing the λ red recombination genes downstream of an arabinose inducible promoter. Subsequently, recombination was carried out as described [62]. Successful mutants were selected by plating on kanamycin (25 *µ*g/ml) and pKD46 (**Table S2**) was removed by growing the plates overnight at 37°C, and then were confirmed with PCR (*mgrB::kanR*, AZ66, **Table S1, S3**). To remove the kanamycin cassette (*ΔmgrB*, AZ132, **Table S1**), pFlp3 containing FLP recombinase and a tetracycline resistance cassette (**Table S2**) was electroporated into AZ66 (**Table S1**) and successful mutants were selected for tetracycline (10 *µ*g/ml) resistance and kanamycin (25 *µ*g/ml) sensitivity, the mutants were then confirmed by PCR (**Table S3**). The plasmid was removed by growing successful mutants on 5% sucrose plates.

To construct all other mutants listed in **Table S1** in their appropriate genetic background, PCR was carried out using Q5 polymerase (NEB) with genomic DNA as template from their respective mutant in MKP103 background [91] and primers listed in **Table S3**, with approximately 500 bp homology on either end of the transposon cassette. λ Red mutagenesis was carried out as described above, and mutants isolated on agar plates containing chloramphenicol (50 *µ*g/ml). Single colonies were purified and verified through PCR. Removal of the chloramphenicol cassette was done as previously described using the plasmid pCre2 [63]. The chromosomal complement of *ΔmgrB* (*mgrB*^*+*^, AZ141, **Table S1**) was constructed as previously described [64] with slight modification. The *mgrB* gene and ∼500 bp upstream and downstream was amplified using WT (AZ55, **Table S1**) genomic DNA as a template. Plasmid pKAS46 (**Table S2**) was digested with NotI and NheI and the *mgrB* PCR product was cloned using the NEBuilder HiFi DNA assembly kit (NEB), followed by transformation into the *E. coli* strain S17-1 λpir. Conjugation with the *ΔmgrB* (AZ132, **Table S1**) and subsequent selection of the complemented strain were carried out as previously described [92].

### Mouse infections and bacterial shedding

C57BL/6J mice were bred and maintained in the animal facility at Biotech Place, Wake Forest Baptist Medical center. Specific Pathogen Free mice (SPF) were obtained from Jackson Laboratory (Bar Harbor, ME). Mice were infected as described [39]. Briefly, 5 to 7 week old mice had food and water removed for 4 hours and orally fed two 50 *µ*l doses an hour apart of ∼10^6^ CFU/100 µl of *K. pneumoniae* suspended in PBS+2% sucrose from a pipette tip.

Fecal collection and quantification of bacterial shedding was done as described previously [39]. To enumerate the bacterial shedding, fecal pellets were homogenized in PBS and serially diluted and plated on selective antibiotic plates. To induce super shedder phenotype mice were infected as described above, and 5 days post-infection administered a single dose of streptomycin (5 mg/ 200 *µ*l in sterile PBS) via oral gavage. *Kpn* was enumerated from fecal shedding for an additional 5 days.

Infections and fecal shedding collection for competition studies (Competition Index [CI]) were conducted as described above with the infection dose containing a 1:1 ratio of MgrB^-^ (AZ132, **Table S1**) and an apramycin resistant derivative of KPPR1 (AZ94, **Table S1**) [39]. For the enumeration of colonization in the GI tract, the cecum, ileum, and colon were removed from mice following CO_2_ (2 liters/min for 5min) euthanasia and subsequent cardiac puncture. The organs were weighed and placed in screw-cap tubes (Fisherbrand; 02-682-558) with 2-3 glass beads (BioSpec Products; 11079127). The organs were diluted 1:10 (weight to volume) in PBS, homogenized, and plated as described above. The CI was calculated using the following formula:

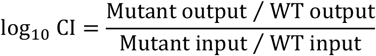

For the transmission studies, 5 mice were separated into individual housing and their water was supplemented with 200 mg/L of streptomycin. Twenty-four hours post-antibiotic exposure, a single mouse was orally infected with either the WT, MgrB^+^, or MgrB^-^ strain. Fecal shedding from the infected index mouse was collected the following day, and colonies enumerated to ensure colonization. Forty-eight hours after infection of the index mouse, 4 uninfected (contact), individually housed mice were placed in the index mouse’s cage one at a time for 1 hour, after which they were returned to their individual cages. Feces was collected for bacterial enumeration from all 5 mice the next day before repeating the exposure. Contact mice were exposed to the index mouse 5 times.

### Growth curves and generation time

Selected *K. pneumoniae* strains were grown overnight at 37°C to stationary phase in LB-Lennox broth (Fisher Scientific, BP1427-2) with constant agitation. The strains were diluted 1:100 in fresh media, and a sample of the subculture was serially diluted and plated onto antibiotic plates for enumeration. The culture was then grown at 37°C with constant agitation with samples collected every hour for 8 hours, serially diluted, and plated for enumeration. For determination of generation time, strains were grown as above until the samples reached an OD_600_ of 0.2; a sample was removed every 15 minutes for up to 75 minutes, diluted and plated for enumeration. The generation time was calculated using the formula G=t/n, where G is the generation time, t is the time interval, and n is the number of generations as calculated by the formula n=3.3log(b/B) where b is the amount of bacteria at the end of a time interval and B is the amount at the beginning of the time interval.

### Lipid A analysis

Lipid A analysis was conducted using the methods described in *Sorensen et al* [65]. Briefly, pelleted bacteria was smeared on a MALDI plate with a sterile inoculation loop. 1 µl of buffer, consisting of 0.2M anhydrous citric acid and 0.1M trisodium citrate dihydrate, was spotted atop the spotted colony. Plate was then incubated at 110°C for 30 minutes to extract membrane lipids. Plate was then rinsed with endotoxin-free water to wash cell debris. 1 µl of 10 mg/mL norharmane matrix suspended in 2:1 chloroform-methanol was spotted onto extracted sample on MALDI plate. Mass spectra was collected using a Bruker Microflex LRF MALDI-TOF MS in negative ion and reflectron mode.

### Fatty acid analysis

The bacterial cell pellet was incubated at 70°C for 1 hour in 500 μl of 90% phenol and 500 μl of water. Samples were then cooled on ice for 5 minutes and centrifuged at 10,000 rpm for 10 minutes. The aqueous layer was collected and 500 μl of water was added to the lower (organic) layer and incubated again. This process was repeated twice, and all aqueous layers were pooled. Two ml of ethyl ether was added to the harvested aqueous layers, this mixture was then vortexed and centrifuged at 3,000 rpm for 5 minutes. The organic phase was then collected, and 2 ml of ether were added back remaining aqueous phase. This process was carried out twice more. The collected organic layer was then frozen and lyophilized overnight. LPS fatty acids were converted to fatty methyl esters, in the presence of 10 μg pentadecanoic acid (Sigma-Aldrich) as an internal standard, with 2 M methanolic HCl (Sigma-Aldrich) at 90°C for 18 hours. The resultant fatty acid methyl esters were analyzed and quantitated by gas chromatography – mass spectrometry (GC-MS) as follows.

GC-MS separation of derivatives was carried out on Shimadzu GC-MS 2010 with split injection with an injection temperature of 280°C. The initial oven temperature was 80°C and held for 1 minute. The temperature was increased 25°C/minute until the oven reached 160°C and held for 1 minute. The temperature was increased 10°C /minute until 265°Cand then increased 1°C/minute until 270°C. The temperature was held at 270°C for 1 minute followed by a 10°C/minute increase to 295°C where it was maintained for 1 minute. Total run time per sample was 25.2 minutes. Column head pressure was 100kPa with an MSD detector. Transfer line temperature was 180°C and mass range was *m/z* 50–400. Identification of metabolites was performed using the standard National Institute of Standards and Technology NIST 08 standard and Golm Metabolome Database (GMD) mass spectra libraries and by comparison with authentic standards. Data processing was performed using a pipeline in KNIME.

### Solid surface survival

To determine the survivability, strains of interest were grown overnight at 37°C with constant agitation. The overnight cultures were centrifuged at 21,000 × *g* for 25 minutes, washed twice with 1X PBS and re-suspended to final adjusted OD_600_ of 4.0. Twenty microliter aliquots of the bacterial suspensions were spotted onto nitrocellulose membranes (MF-Millipore, HAWP02500) on 2-ml pads of 1%agarose (Fisher, BP160-500) in 6-well polystyrene plates (Costar, 3516) and air dried. At designated timed intervals the membrane was removed, placed in 1 ml PBS, vortexed to dislodge the bacteria, and the suspension serially diluted and plated onto LB agar plates (Fisher, BP9745-2) In between sampling the plates were sealed with parafilm (Bemis, PM-999) and placed in the dark at room temperature.

### Biofilm quantification

For biofilm assays overnight cultures were grown as described above in LB-Lennox broth and diluted 1:100 in M63 media with 0.5% glucose. One-hundred microliters of the bacterial suspension was placed in the wells of a 96-well round-bottom culture plate (Costar), sealed (Fisherbrand Thermal Adhesive Sealing Film), and incubated for 24 hours at 37° C. Staining and quantification were carried out as described [66].

### Mucoviscosity assay

Mucoviscosity of the *Kpn* isolates was determined using a low speed centrifugation (1000 x *g*, 5 min) of bacterial cultures adjusted to 1 OD_600_ and measuring the OD_600_ of the supernatant as previously described [67].

### Mucin binding assay

Mucin binding was determined as previously described [45]. Nunc Polysorp plates (Thermo Fisher Scientific, 475094) were coated with 100 *µ*l of 50 *µ*g semi-purified Millipore mucin (Cat. 499643-500MG) by centrifugation (250 x *g*, 3 min) and incubated overnight at 37°C. The bound mucin was washed 3 times, and then ∼10^5^ CFU of *Kpn* in 100*µ*l was added. The plate was centrifuged (250 x *g*, 3 min) and incubated at 37°C for 60 min. The wells were washed 15 times to remove any unbound bacteria. Subsequently the wells were treated with 200 *µ*l of PBS-0.5% Triton X-100 (Sigma-Aldrich) for 30 minutes followed by vigorous mixing. Samples were then plated in triplicate from the wells for enumeration.

### Antimicrobial resistance assays

The polymyxin and lysozyme resistance assays were carried out as described [68] [69]. For the lysozyme assay *Kpn* strains were grown overnight then diluted 1:100 in fresh media and grown to mid-log phase (OD_600_ 0.4-0.5), centrifuged (21,000 x *g*, 20 min) and washed with PBS twice before being diluted 1:100 in Tris-EDTA (TE) buffer with 1% Tryptic Soy Broth (TSB) (BD). Ninety-six well plates were prepared with 100 *µ*l of appropriate concentrations of chicken egg lysozyme suspended in TE + 1% TSB and 100 *µ*l of the bacterial suspension was added to wells in triplicate. The plate was incubated at 37°C for 1 hour before samples were taken from each well, diluted, and plated for enumeration.

### Gel electrophoresis and analysis

Western-blots were carried out as previously described [70] with modifications. Bacterial cultures were grown overnight in cation adjusted Mueller Hinton Broth (caMHB), diluted 1:100 in fresh caMHB and grown for 16 hours to stationary phase. 500 *µ*l of bacterial culture was pelleted (21,000 x *g*, 25min), re-suspended in a volume of 2x Laemmli sample buffer (BioRad) equal to 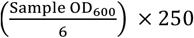. Samples were boiled at 100°C for 20 minutes. Fifteen microliters of each sample were loaded into the wells of a 12% polyacrylamide gel (Bio-Rad, 4561045) and run at 150 V for 75 min. Transfer from the gel to a nitrocellulose membrane (Bio-Rad, 1704158) was carried out using the Bio-Rad Trans-Blot Turbo Transfer System. The membrane was probed with anti-RNA Sigma S Antibody (BioLegend, Cat. 663703) and subsequently with Goat Anti-Mouse IgG (H + L)-HRP Conjugate (Bio-Rad, Cat. 1706516). The blot was developed with Clarity Western ECL Substrate (Bio-Rad, 170-5061). Final imaging was done using the Bio-Rad ChemiDoc MP imaging system.

### Statistical analysis

All statistical analyses were performed using GraphPad Prism 9.0 (GraphPad Software, Inc., San Diego, CA). Unless otherwise specified, differences were determined using the Mann-Whitney *U* test (comparing two groups), the Kruskal-Wallis test with Dunn’s post-analysis (comparing multiple groups), the Wilcoxon sign ranked test, or Fisher’s exact test.

## Figure legends

**Fig. S1**. Growth and generation time are not impacted by MgrB inactivation. (***A***) Growth curves and (***B***) generation times in nutrient rich LB media. (***C***) Growth curves and (***D***) generation times in M63 minimal media supplemented with 0.5% fucose, an alternative carbon source present in the GI-tract that can be metabolized by *Kpn*.

**Fig. S2**. GI colonization defect due to MgrB inactivation is conserved across strains. Mice were orally infected with either the ST1322 or the *ΔmgrB* isogenic mutant, and feces was collected on the days indicated. Each point indicates a single mouse on a given day, the bars indicate the median shedding, and the dotted line indicates the limit of detection, significance shown is between WT and MgrB^-^. Statistical significance was determined with Mann-Whitney U tests at each time point. * *P* < 0.05, ** *P* < 0.005, n.s, not significant.

**Fig. S3**. MgrB inactivation does not impact (***A***) biofilm formation or (***B***) hypermucoviscosity.

**Fig. S4**. Shedding of (***A***) index and (***B***) contact mice through transmission studies.

**Fig. S5**. RpoS protein levels in stationary phase cultures of WT, MgrB^-^, and MgrB^+^ grown in cation adjusted Mueller Hinton Broth through Western-blot analysis. Shown is a representation of three independent experiments.

**Table S1.**
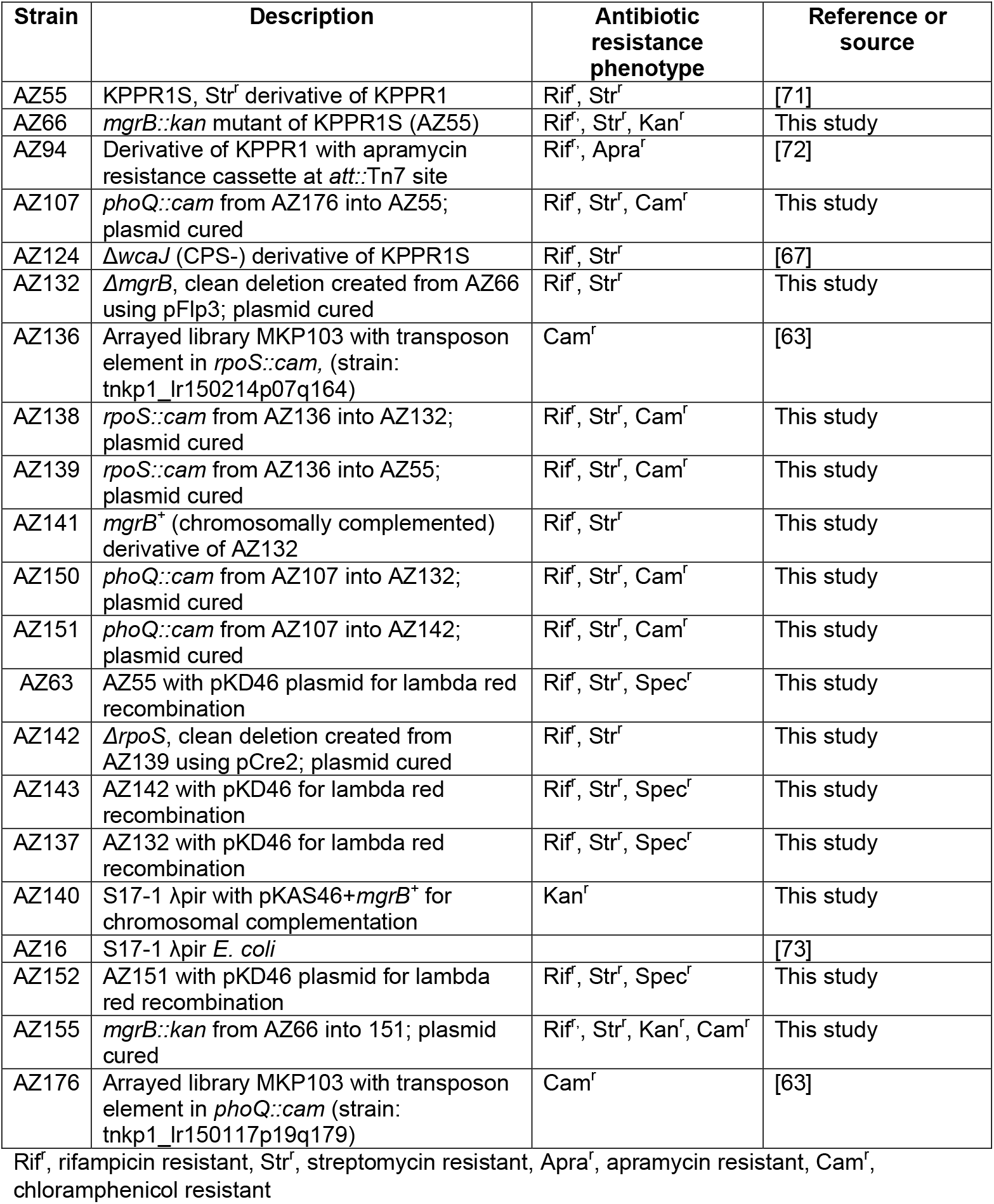

**Table S2.**
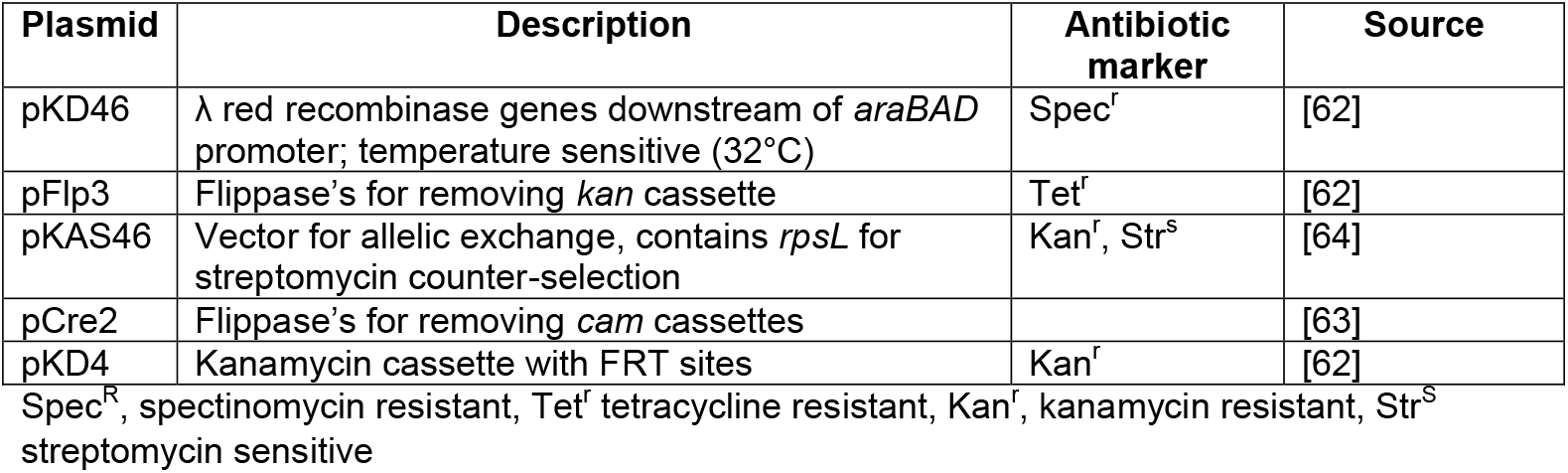

**Table S3.**
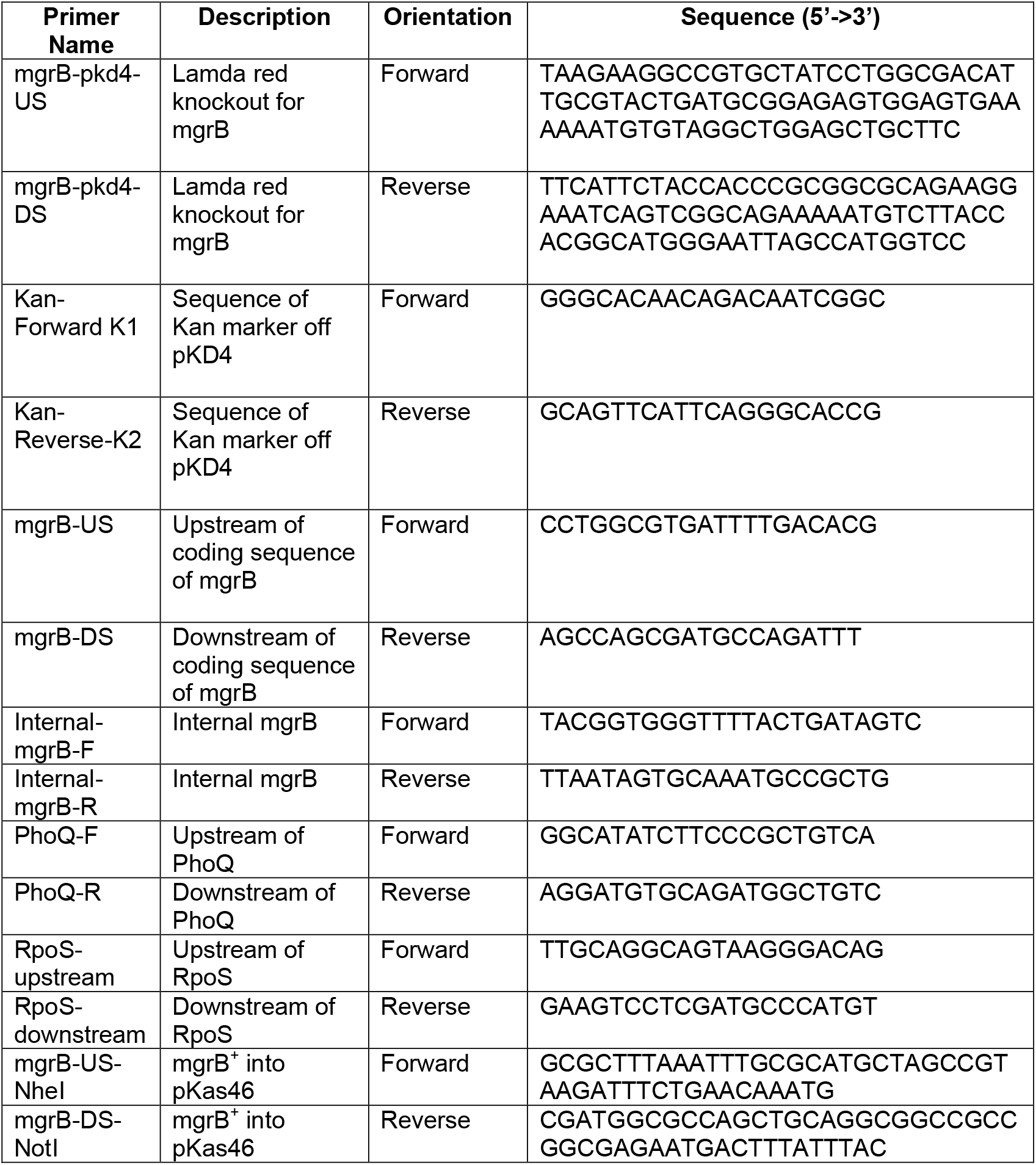

